# Mechanical Ventilation Impairs Glymphatic Function and Induces Delirium-Like Behavior in Mice

**DOI:** 10.64898/2025.12.12.694005

**Authors:** Guojun Liu, Jianbo Wang, Tianyi Tong, SiRu Liang, Evan Newbold, Erik Kroesbergen, Sungha Hong, Antonio Ladrón-de-Guevara, Hans Christian Boesen, Maiken Nedergaard, Ting Du

**Affiliations:** Center for Translational Neuromedicine, Department of Neurosurgery, University of Rochester Medical Center, Rochester, NY, 14642, USA; Department of Pharmacology and Physiology, University of Rochester Medical Center, Rochester, NY 14642, USA; Department of Brain and Cognitive Sciences, University of Rochester, NY 14642, USA; Intensive Care Unit, Department of Anesthesiology, Zealand University Hospital, 4600, Køge, Denmark; Center for Translational Neuromedicine, Faculty of Health and Medical Sciences, University of Copenhagen, 2200, Copenhagen, Denmark

**Keywords:** Delirium, Mechanical ventilation, Glymphatic transport, CSF efflux, Glymphatic clearance, Cytokines

## Abstract

Delirium is a common and serious complication in critically ill patients, arising from multiple overlapping risk factors. Its pathogenesis remains poorly understood because it reflects the combined effects of interacting clinical insults, and because appropriate experimental models for mechanistic studies have been lacking. Delirium most commonly develops in patients undergoing major surgery or requiring intensive care, where inhalational anesthesia, opioid sedation, and mechanical ventilation are common clinical exposures.

Here, we tested the hypothesis that these clinically relevant interventions are sufficient to induce delirium-like behavior in healthy young wild-type mice by disrupting cerebrospinal fluid (CSF) glymphatic transport. The analysis showed that inhalational anesthesia combined with mechanical ventilation increased intracranial pressure (ICP) and induced a sustained suppression and rerouting of glymphatic waste clearance, resulting in impaired waste clearance, and acute cytokine accumulation.

These findings identify disruption of brain fluid dynamics as a potential mechanism contributing to delirium in perioperative and intensive care settings. Equally important, this model represents a distinct experimental platform that combines clinically relevant mechanical ventilation, inhalational anesthesia, and opioid analgesia to produce delirium-like behavioral abnormalities in otherwise healthy mice. It therefore provides a powerful tool for future mechanistic studies of delirium pathogenesis.

## INTRODUCTION

Delirium is an acute neuropsychiatric syndrome characterized by acute and fluctuating disturbances in attention, awareness, cognition, and sleep that cannot be explained by pre-existing neurodegenerative disease or another acute neurological disorder^1^. Delirium occurs in ∼20–50% of ICU patients, rising to ∼80% among those who require mechanical ventilation ^2,3^. Delirium affects an estimated 1–2 million critically ill patients admitted to intensive care units in the United States each year, making it one of the most common forms of acute brain dysfunction in critical illness ^4^. The risk landscape for ICU delirium is complex and includes advanced age ^5^, pre-existing cognitive impairment ^6,7^, central nervous system (CNS) disease ^2,8^, and systemic comorbidities such as chronic cardiac, hepatic, or renal dysfunction (e.g., chronic heart, liver, or kidney disease) ^9–11^. ICU exposures, including inhalation anesthetics, sedatives such as benzodiazepines or propofol, opioid analgesia, mechanical ventilation, and sleep deprivation^1,2,7,12,13^, further increases delirium risk. Because these factors often occur simultaneously in critically ill patients, it remains difficult to determine how individual exposures, and their associated physiological perturbations interact to precipitate acute brain dysfunction.

We reasoned that glymphatic failure may represent a convergent mechanism linking ICU exposures to delirium. The glymphatic system is a brain-wide perivascular fluid-transport pathway in which cerebrospinal fluid (CSF) enters along peri-arterial spaces, exchanges with interstitial fluid (ISF) within the parenchyma, and clears solutes along peri-venous routes toward meningeal lymphatics and cervical lymph nodes ^14–16^. Glymphatic flux is strongly modulated by the sleep–wake state ^17,18^, anesthetics ^19,20^, aging ^21^, intracranial pressure (ICP) ^22^ and respiratory dynamics ^23^. Many ICU stressors – sleep deprivation, deep sedation/anesthesia, systemic inflammation/infection, and mechanical ventilation – are known or suspected to suppress glymphatic function ^24–26^. We therefore hypothesized that mechanical ventilation-associated glymphatic dysfunction may reduce brain solute clearance, thereby increasing the local availability of inflammatory mediators and other bioactive metabolites that could contribute to acute brain dysfunction ^27^.

A major barrier to testing this idea is the limited availability of translational experimental models that isolate the contributions of clinically relevant ICU exposures while capturing the combined respiratory, anesthetic, sedative, and physiological perturbations experienced by mechanically ventilated patients. Animal studies of delirium have modeled systemic inflammation, urinary tract infection, bone fracture, sleep deprivation, administration of anesthetics, and metabolic stress ^28,29^. Although these approaches have provided mechanistic insight, they do not fully reproduce the concurrent respiratory, anesthetic, sedative, and physiological perturbations that accompany prolonged exposure to inhalation anesthesia, fentanyl administration and mechanical ventilation in the ICU.

Here, we developed a murine model incorporating key features of intensive care exposure, including mechanical ventilation, inhalation anesthesia, and opioid analgesia. We used this model to determine whether a brief combined ICU-relevant exposure is sufficient to produce delirium-like behavioral changes detectable late - 24 hrs - in otherwise healthy mice. Using this model, we demonstrate that mechanical ventilation acutely disrupts brain fluid homeostasis by impairing glymphatic transport, altering CSF efflux pathways, reducing brain-to-blood clearance, and eliciting a brain-predominant cytokine response.

## RESULTS

### Delirium-like behavior develops after mechanical ventilation

To determine whether mechanical ventilation (MV) elicits behavioral changes, mice in the MV group underwent endotracheal intubation and received 2.5% isoflurane through the ventilator circuit during 2 h of mechanical ventilation, with adjunct intraperitoneal fentanyl. Duration-matched control mice received 2.5% isoflurane via a nose cone (NC) with adjunct intraperitoneal fentanyl, but were not intubated or mechanically ventilated; unexposed mice served as Control animals. During mechanical ventilation, end-tidal CO₂ was continuously monitored and maintained within the physiological range, oxygenation was monitored using a sensor affixed to the shaved hindlimb, and normothermia was maintained using a 37°C heating pad. After 2 h of MV or NC, mice were returned to their home cages; 24 h later, behavior was assessed (**Fig. 1a**).

**Figure 1.**
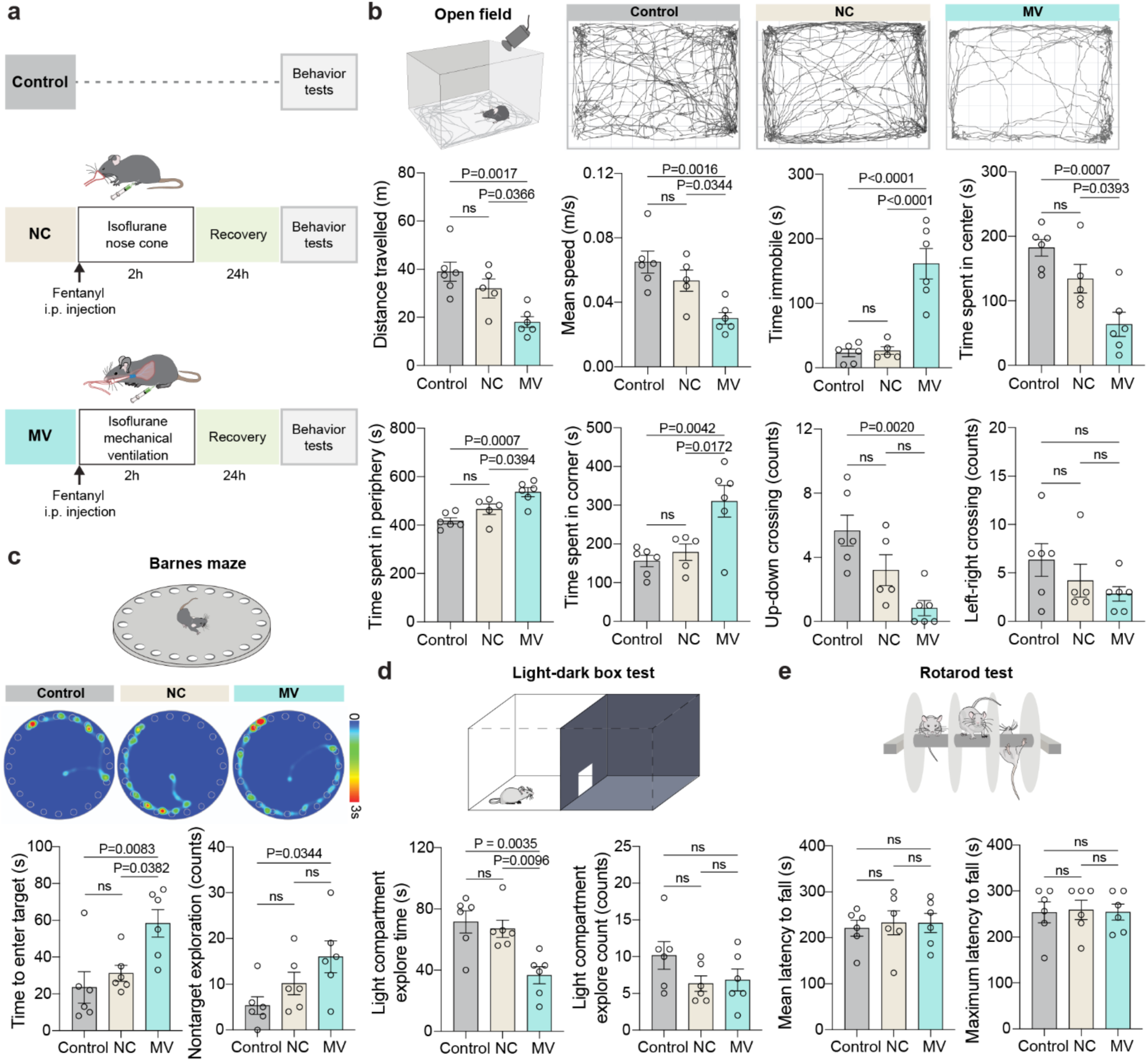
Mechanical ventilation induces delirium-like behavioral abnormalities 24h after exposure in mice. **(a)** Experimental timeline. Control mice were tested at the matched time point without anesthesia or ventilation. Mice in the nose cone (NC) group received 2 h of 2.5% isoflurane via nose cone, whereas mice in the mechanical ventilation (MV) group underwent 2 h of mechanical ventilation under 2.5% isoflurane anesthesia. NC and MV mice received intraperitoneal (i.p.) fentanyl before anesthesia. Behavioral tests were conducted 24 h later. **(b)** Representative open-field trajectories from control, NC, and MV mice during a 10-min open-field test. Bar graphs show open-field locomotor activity, quantified as total distance traveled (m), mean speed (m/s), and immobility time (s); spatial distribution, quantified as time spent in the center, periphery, and corners of the arena (s); and zone transitions, quantified as up–down and left–right crossings (counts) during the 10-min session. **(c)** Representative Barnes maze occupancy heatmaps and trajectory plots from Control, NC, and MV mice. Bar graphs show Barnes maze performance, quantified as time to enter the target hole (s) and non-target hole explorations (counts). **(d)** Quantification of light–dark box performance, including light compartment entries (counts) and time spent in the light compartment (s). **(e)** Bar graphs showing rotarod performance, quantified as mean latency to fall (s) and maximum latency to fall (s). Data are presented as mean ± SEM. One-way ANOVA followed by Tukey’s multiple-comparisons test was performed. n=5-6 mice per group. ns, not significant.

In the open field test, MV-exposed mice showed reduced locomotor activity and increased thigmotaxis relative to Control (**Fig. 1b**): total distance and mean speed were reduced, immobility time increased, time in center decreased with reciprocal increases in periphery/corner dwell, and zone crossings (up– down) were markedly reduced. NC mice did not differ from Control. These findings indicate diminished exploration and increased avoidance/anxiety-like behavior 24 h after MV, but not after matched NC exposure. In the Barnes maze, MV mice took longer to enter the target and made more non-target entries (**Fig. 1c**), indicating a slower, less efficient, and more error-prone search strategy. In the light–dark box test, MV mice showed a significant reduction in time spent exploring the light compartment, while the number of light-compartment entries was not significantly changed (**Fig. 1d**). This pattern indicates reduced sustained exploration of the light compartment despite preserved entry frequency, consistent with increased light avoidance. To determine whether the reduced exploratory behavior observed after mechanical ventilation could be attributed to gross motor impairment, we performed the rotarod test (**Fig. 1e**). MV mice showed no significant differences in mean or maximum latency to fall compared with Control or NC mice, suggesting preserved motor coordination and balance. These findings indicate that the behavioral alterations observed in the open field and light– dark box tests are unlikely to be explained solely by gross motor dysfunction.

Collectively, these behavioral data indicate that MV, but not matched NC exposure, induced a constellation of behavioral alterations consistent with delirium-like dysfunction, increased anxiety-like avoidance, and altered spatial search behavior, without evidence of gross motor impairment.

### Mechanical ventilation induces acute glymphatic dysfunction that persists into early recovery

Because behavioral alterations were evident 24 h after mechanical ventilation, we next asked whether mechanical ventilation produces glymphatic dysfunction that persists beyond the ventilation period itself. To test this, mice underwent 2 h of MV or nose-cone anesthesia (NC) under the same isoflurane/fentanyl regimen, were allowed to recover awake for 1 h, and were then re-anesthetized with ketamine/xylazine (KX) for glymphatic imaging. BSA–Alexa Fluor 647 (BSA-647) was injected into the cisterna magna (CM), followed by 30 min of transcranial macroscopic imaging and subsequent ex vivo brain analysis (**Fig. 2a**).

**Figure 2.**
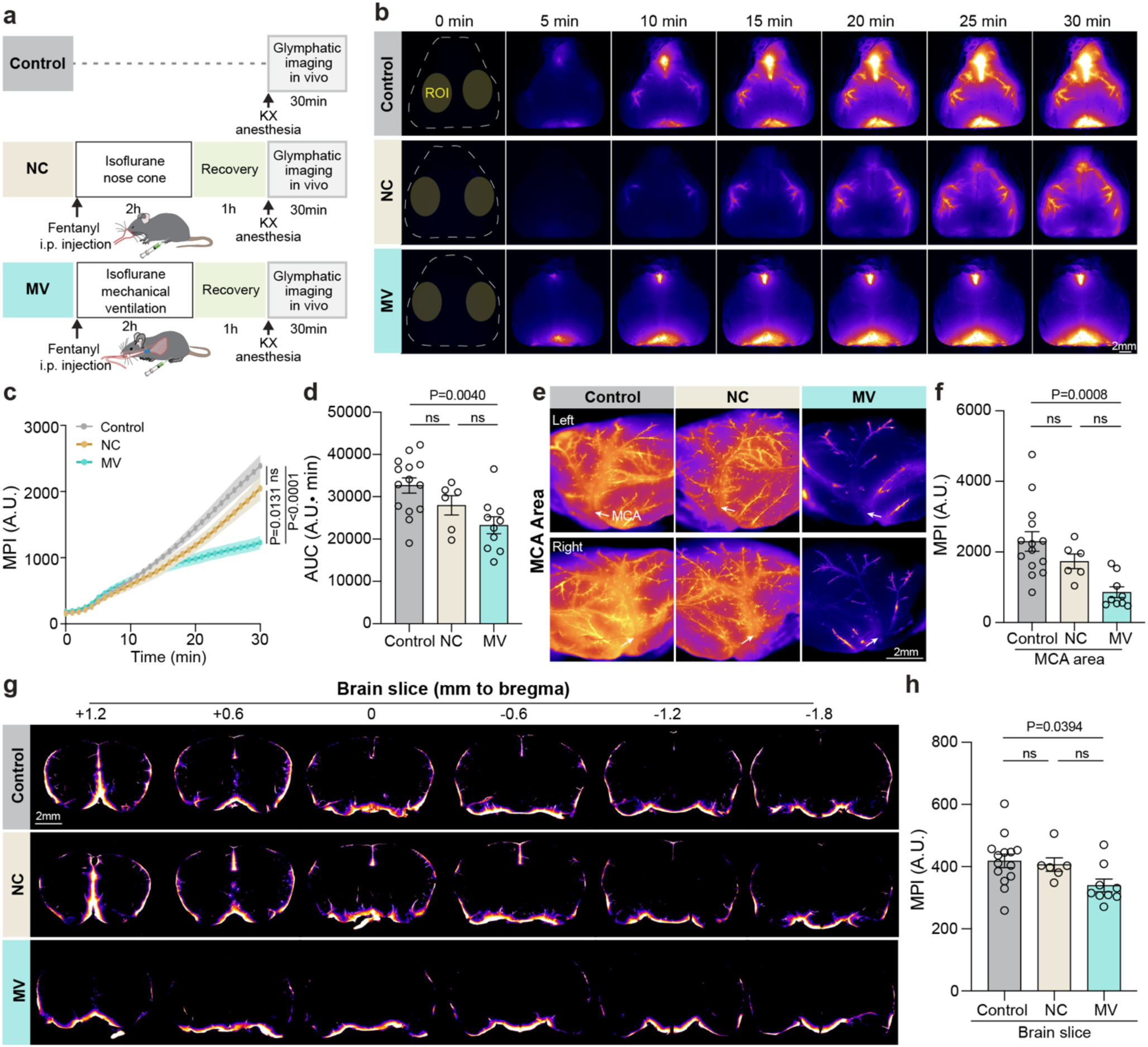
Glymphatic transport is suppressed after 1 h recovery from mechanical ventilation. **(a)** Experimental timeline for assessing glymphatic influx in vivo. Control mice received no prior nose-cone (NC) or mechanical ventilation (MV) exposure and were anesthetized with ketamine/xylazine (KX) for cisterna magna (CM) injection of BSA–Alexa Fluor 647 (BSA-647), followed by 30 min of transcranial imaging. NC mice received 2 h of 2.5% isoflurane anesthesia via a nose cone, whereas MV mice underwent 2 h of mechanical ventilation under 2.5% isoflurane anesthesia. NC and MV mice received intraperitoneal (i.p.) fentanyl before isoflurane exposure. After a 1-h awake recovery period, NC and MV mice were re-anesthetized with KX for CM injection of BSA-647, followed by 30 min of transcranial imaging. **(b)** Representative transcranial fluorescence images showing time-lapse intracisternal tracer influx in Control, NC, and MV mice. Scale bar, 2 mm. **(c)** Quantification of mean pixel intensity (MPI) of tracer influx over 0–30 min within the middle cerebral artery (MCA) territory. Statistical annotations indicate group comparisons at the 30-min time point. **(d)** Area under the curve (AUC) for 30-min BSA-647 influx from the data shown in panel c. **(e)** Representative high-resolution ex vivo fluorescence images of intact whole brains showing tracer distribution in the MCA territory. **(f)** Quantification of ex vivo MPI in the MCA territory. **(g)** Representative ex vivo fluorescence images of coronal brain sections from anterior to posterior. Scale bar, 2 mm. **(h)** Quantification of MPI in ex vivo coronal sections. Data are presented as mean ± SEM. Statistical significance was determined by two-way repeated-measures ANOVA with Sidak’s multiple-comparisons test for panel c, and by one-way ANOVA followed by Tukey’s post hoc test for panels d, f, and h. (b–d) n = 14 Control mice, n = 6 NC mice, and n = 10 MV mice. (e–h) n = 14 Control mice, n = 6 NC mice, and n = 9 MV mice.. ns, not significant.

In vivo imaging revealed markedly reduced tracer influx along the middle cerebral artery (MCA) territory in MV mice compared with Control mice (**Fig. 2b and c; Video 1**). In contrast, tracer influx was largely preserved in NC mice, indicating that isoflurane plus fentanyl exposure without mechanical ventilation was not sufficient to reproduce the glymphatic impairment. Quantification of the area under the curve (AUC) confirmed reduced tracer influx in MV mice after 1 h of awake recovery (**Fig. 2d**). We next performed high-resolution ex vivo imaging of whole brains to assess tracer distribution within the MCA territory (**Fig. 2e**). Consistent with the in vivo findings, ex vivo whole-brain imaging showed a pronounced reduction in MCA-associated tracer signal in the MV group compared with Control group (**Fig. 2f**). Serial coronal brain sections further demonstrated reduced BSA-647 distribution across the anterior–posterior extent of the brain in MV mice (**Fig. 2g and h**).

We next asked whether glymphatic impairment was already present during mechanical ventilation, before the recovery period. Because glymphatic influx is strongly influenced by anesthetic state, and is robust under KX anesthesia^17,19^, KX was administered before tracer CM injection and imaging in a separate During MV cohort to reduce differences in anesthetic conditions relative to KX-anesthetized Control mice. In this cohort, mice were mechanically ventilated under isoflurane anesthesia with i.p. fentanyl. At 1.5 h after MV onset, while ventilation was maintained, mice received KX, followed by cisterna magna injection of BSA-647. Transcranial imaging was then performed over the subsequent 30 min while MV continued. The same KX-anesthetized Control group used in Fig. 2 served as a shared reference group (**Supplementary Fig. 1a**). Glymphatic influx was already markedly reduced during MV compared with KX-anesthetized controls (**Supplementary Fig. 1b–d; Video 2**). High-resolution ex vivo imaging of intact whole brains focused on the MCA territory, together with coronal-section imaging, showed the same pattern: reduced tracer accumulation in the MCA territory and reduced tracer distribution across coronal brain sections during MV (**Supplementary Fig. 1e–h**).

Together, these data show that glymphatic tracer influx was impaired during MV and remained impaired after 1 h of awake recovery. This impairment was not observed in matched NC mice exposed to isoflurane plus fentanyl without mechanical ventilation.

### Mechanical ventilation produces time-dependent changes in CSF tracer distribution across downstream efflux compartments

We next asked whether mechanical ventilation altered CSF tracer distribution across downstream efflux compartments during ventilation and early recovery. Using the experimental paradigms shown in Figs. 3a and 4a, and Supplementary Figs. 2a and 3a, tracer distribution was assessed ex vivo in the whole brain, skull compartments, spinal cord, and superficial and deep cervical lymph nodes (scLNs and dcLNs) following in vivo imaging of glymphatic influx.

**Figure 3.**
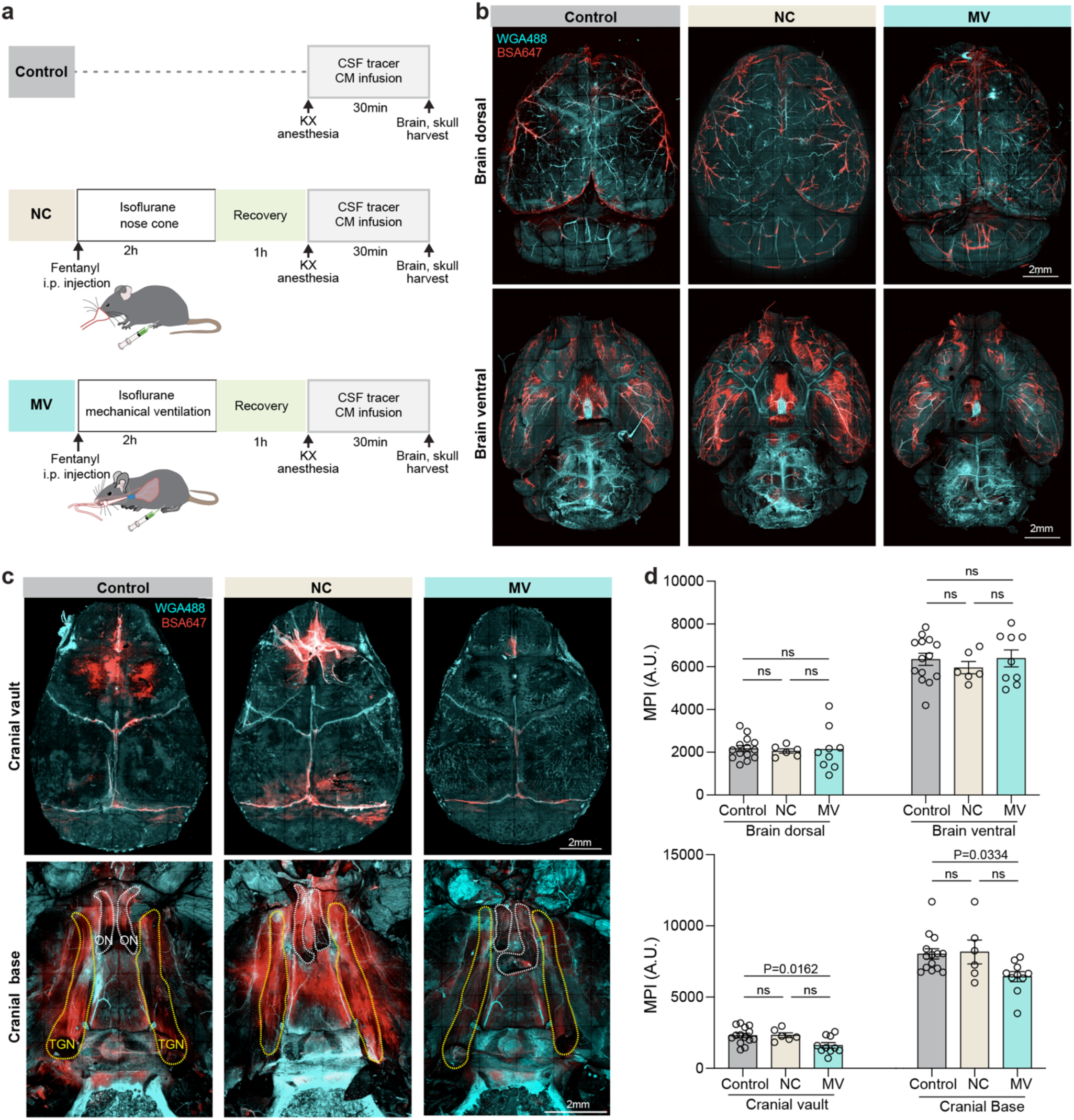
Mechanical ventilation reduces cranial-base CSF tracer distribution after 1 h recovery. **(a)** Experimental timeline for ex vivo assessment of intracisternally delivered tracer distribution. Following cisterna magna (CM) injection of BSA–Alexa Fluor 647 (BSA-647) and a 30-min influx period, as described in Fig. 2a, tissues including the whole brain and skull were harvested from Control, nose cone (NC), and mechanical ventilation (MV) mice and imaged ex vivo by fluorescence microscopy. **(b)** Representative dorsal and ventral whole-brain fluorescence images from Control, NC, and MV mice. Scale bar, 2 mm. **(c)** Representative fluorescence images of the cranial vault and cranial base. Scale bar, 2 mm. **(d)** Quantification of mean pixel intensity (MPI) in the dorsal brain, ventral brain, cranial vault, and cranial-base regions associated with the olfactory nerve (ON) and trigeminal nerve (TGN). Data are presented as mean ± SEM. Statistical significance was determined by one-way ANOVA followed by Tukey’s post hoc test. For dorsal and ventral whole-brain analyses, n = 14 Control mice, n = 6 NC mice, and n = 9 MV mice; for cranial vault and cranial-base analyses, n = 14 Control mice, n = 6 NC mice, and n = 10 MV mice. ns, not significant.

**Figure 4.**
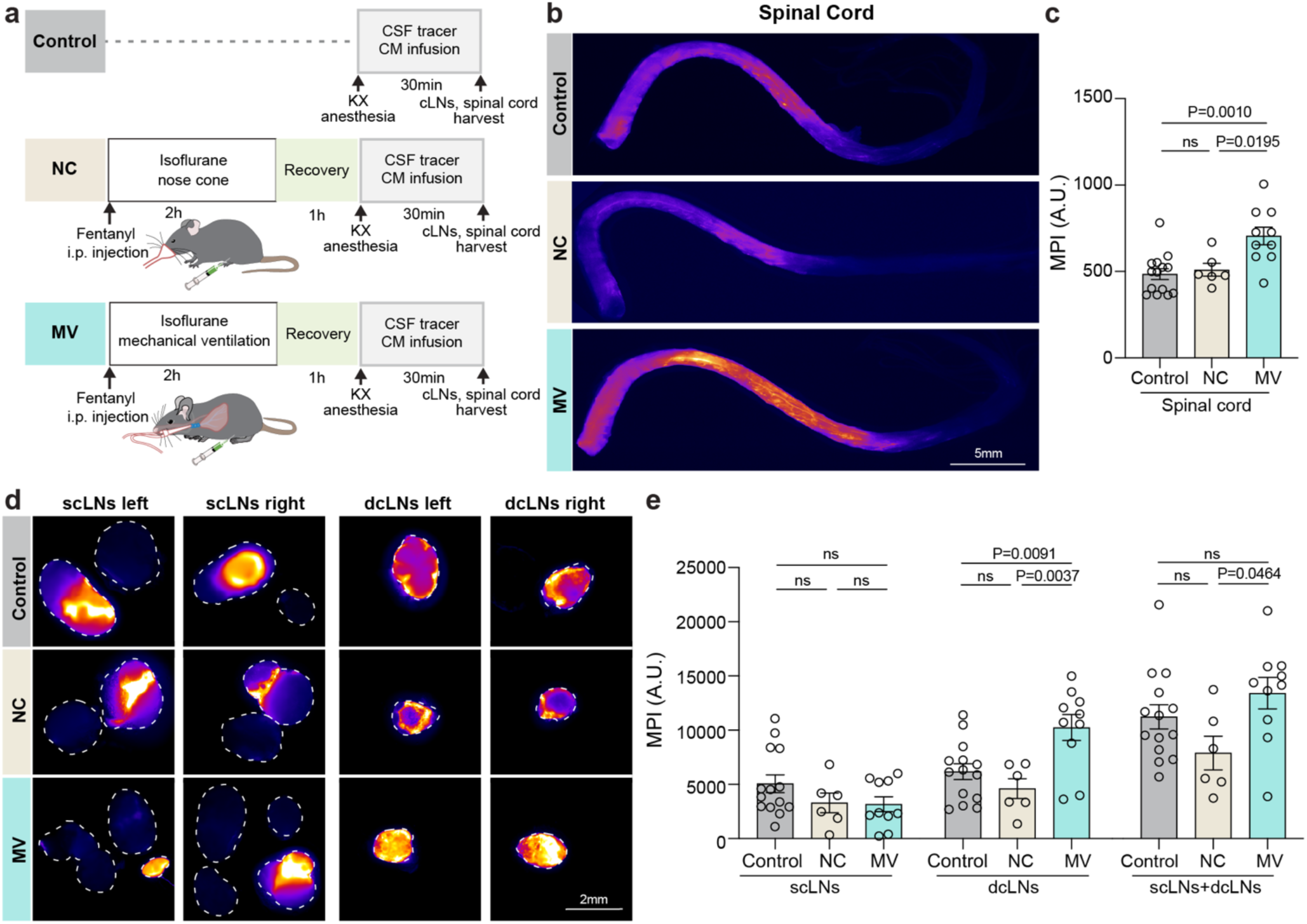
Mechanical ventilation alters CSF tracer efflux pathways after 1 h recovery. **(a)** Experimental timeline for ex vivo assessment of intracisternally delivered tracer distribution. Following cisterna magna (CM) injection of BSA–Alexa Fluor 647 (BSA-647) and a 30-min influx period, as described in Fig. 2a, the tissues including the spinal cord, superficial cervical lymph nodes (scLNs), and deep cervical lymph nodes (dcLNs) were harvested from Control, nose cone (NC), and mechanical ventilation (MV) mice and imaged ex vivo by fluorescence microscopy. **(b)** Representative fluorescence images of the spinal cord. Scale bar, 5 mm. **(c)** Bar graph shows quantification of spinal cord MPI. **(d)** Representative fluorescence images of scLNs and dcLNs from Control, NC, and MV mice. Scale bar, 2 mm. **(e)** Bar graphs show quantification of MPI in scLNs, dcLNs, and combined scLNs + dcLNs. Data are presented as mean ± SEM. Statistical significance was determined by one-way ANOVA followed by Tukey’s post hoc test for panels c and e. n = 14 Control mice, n = 6 NC mice, and n = 10 MV mice. ns, not significant.

During MV, tracer distribution was reduced in the dorsal and ventral brain, cranial vault, and cranial-base regions associated with the olfactory nerve (ON) and trigeminal nerve (TGN) (**Supplementary Fig. 2b and c**), indicating a broad acute disruption of brain and skull tracer distribution during ventilation. We next examined whether this pattern persisted after 1 h of recovery from MV. One hour after recovery, tracer intensity in the dorsal and ventral brain was comparable among Control, NC, and MV mice (**Fig. 3b and d**). However, tracer signal remained reduced in the cranial vault and ON- and TGN-associated cranial-base regions in MV mice (**Fig. 3c and d**). These findings indicate that the broad reduction in tracer distribution observed during MV partially resolves during early recovery but persists in cranial vault and cranial-base regions.

Peripheral CSF tracer distribution showed a distinct time-dependent response. During MV, spinal cord tracer accumulation was reduced, whereas tracer accumulation in scLNs was increased and dcLNs were unchanged (**Supplementary Fig. 3**). By 1 h after recovery, this pattern shifted: spinal cord tracer accumulation increased (**Fig. 4b and c**), scLN tracer levels were no longer elevated, and dcLN tracer accumulation increased in MV mice, while NC mice showed no significant changes (**Fig. 4d and e**).

Importantly, the combined tracer signal in scLNs and dcLNs did not differ from that in Control mice at either time point, indicating that MV altered the distribution of tracer between superficial and deep cervical lymphatic pathways rather than the overall amount of tracer recovered across the cervical lymph nodes.

Together, these findings show that the broad reduction in intracranial tracer distribution during MV partially resolved during early recovery, whereas persistent reductions in cranial vault and cranial-base tracer signal were accompanied by time-dependent changes in spinal and cervical lymph-node tracer distribution.

### Mechanical ventilation acutely elevates ICP

Given the role of ICP in shaping CSF egress^30,31^, we next monitored ICP continuously during 2 h of mechanical ventilation (MV) and the subsequent 1 h recovery period and compared ICP with that in NC-mice exposed to matched isoflurane and intraperitoneal (i.p.) fentanyl without ventilation (**Fig. 5a**). End-tidal CO₂ was maintained within the physiological range throughout MV. During the 2 h ventilation period, ICP was significantly elevated in MV mice compared with NC mice (**Fig. 5b**).

**Figure 5.**
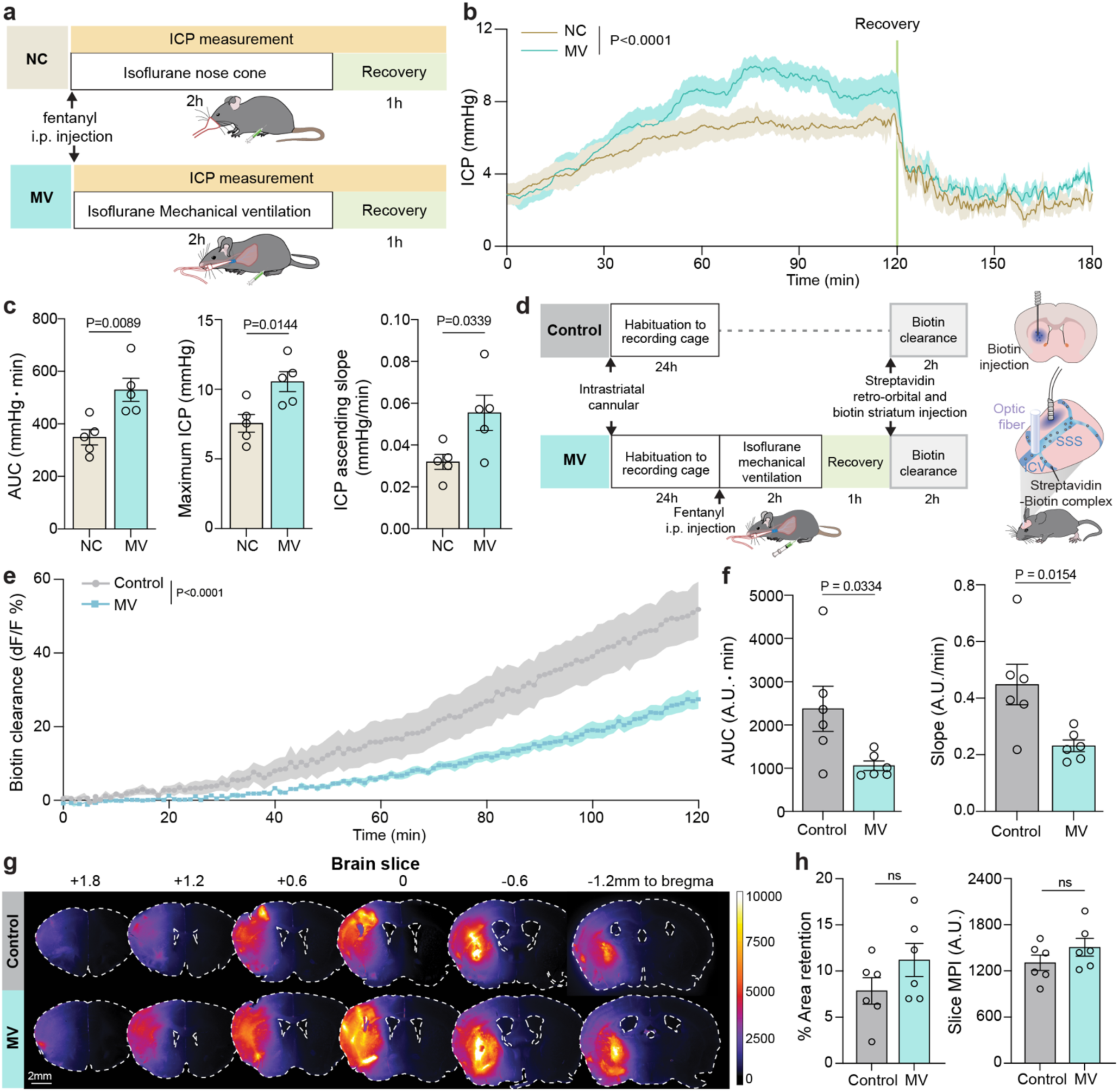
Mechanical ventilation suppresses glymphatic clearance. **(a)** Experimental timeline. Intracranial pressure (ICP) was recorded continuously during 2 h of 2.5% isoflurane anesthesia delivered via a nose cone (NC) or during mechanical ventilation (MV) under 2.5% isoflurane anesthesia, with intraperitoneal (i.p.) fentanyl administered before isoflurane exposure, followed by 1 h of recovery. **(b)** Time course of ICP during NC or MV exposure. **(c)** Summary quantification of ICP parameters derived from panel b, including area under the curve (AUC) over the 120-min exposure period, maximum ICP, and the rising slope of ICP (mmHg/min). (d) Assay schematic for interstitial fluid (ISF)-to-plasma clearance after MV. Mice underwent 2 h of MV under 2.5% isoflurane anesthesia with i.p. fentanyl. After a 1-h awake recovery period, streptavidin was delivered into the bloodstream via retro-orbital injection, and Cy5-biotin was injected into the striatum. Cy5-biotin cleared from brain ISF to plasma, where it bound circulating streptavidin and was retained in the circulation for more than 2 h, enabling stable quantification of brain-to-plasma clearance. An optic fiber placed over the confluence of the superior sagittal sinus (SSS) and inferior cerebral vein (ICV) detected the biotin–streptavidin efflux signal. (e) Time course of SSS-ICV confluence fluorescence over 2 h, expressed as ΔF/F₀%. (f) Summary quantification of fluorescence dynamics derived from panel e, including AUC and the slope of signal increase. (g) Representative ex vivo coronal brain sections from anterior to posterior showing brain retention of Cy5-biotin after intrastriatal Cy5-biotin and retro-orbital streptavidin injections. Scale bar, 2 mm. (h) Quantification of brain retention as percent fluorescent area and mean pixel intensity (MPI). Data are presented as mean ± SEM. Statistical significance was determined by two-way repeated-measures ANOVA for panels b and e, and by two-tailed unpaired t-test for panels c, f, and h. n = 5 mice per group for panels b and c; n = 6 mice per group for panels e–h. ns, not significant.

Quantification of the ICP AUC over the 120-min exposure period showed an approximately 51.8% increase in MV mice compared with NC mice (**Fig. 5c**). The maximum ICP reached during the 2 h period was also significantly higher in MV mice than in NC mice (NC: 7.56 ± 0.63 mmHg; MV: 10.55 ± 0.72 mmHg; P = 0.0144; Fig. 5c). In addition, ICP rose more steeply during MV than during NC anesthesia (**Fig. 5c**), indicating that the MV condition elevates ICP beyond the effect of matched isoflurane and opioid exposure alone. This ICP elevation occurred during the ventilation period, coincident with the broad reduction in CSF tracer distribution detected during ongoing MV.

### Mechanical ventilation impairs awake brain-to-blood tracer clearance

A limitation of the glymphatic influx experiments described above is that CSF tracer delivery and macroscopic imaging were performed under KX anesthesia. We therefore asked whether prior MV exposure altered an awake, freely moving measure of brain-to-blood clearance of an intraparenchymally delivered tracer rather than CSF tracer influx^32^.

To determine whether MV altered this awake brain-to-blood tracer clearance readout, mice were first implanted with an intra-striatal cannula and habituated to the recording cage for 24 h before data collection to minimize stress-related confounding. MV mice then received i.p. fentanyl and underwent 2 h of isoflurane mechanical ventilation, followed by 1 h of awake recovery before the brain clearance assay; Control mice underwent the same habituation and clearance procedure without MV exposure (**Fig. 5d**).

For this assay, Cy5-conjugated biotin tracer (Cy5-biotin) was injected into the striatum through a preimplanted cannula, and streptavidin was administered intravenously by retro-orbital injection. Once Cy5-biotin entered the bloodstream, it bound circulating streptavidin, forming a stable complex that enabled cumulative detection of vascular Cy5 signal over time. An optical fiber positioned over the confluence of the superior sagittal sinus (SSS) and inferior cerebral vein (ICV) recorded the vascular fluorescence signal using fiber photometry (**Fig. 5d**). Mice were habituated to the recording cage for 24 h before data collection to minimize stress-related confounding.

The vascular fluorescence signal at the SSS–ICV confluence, used here as a readout of brain-to-blood tracer clearance, was significantly blunted in mechanically ventilated mice compared with controls (P < 0.0001; **Fig. 5e**). Quantification of both the AUC and the signal slope confirmed a marked reduction in vascular tracer signal after MV (**Fig. 5f**). Quantification of residual Cy5-biotin in ex vivo brain sections revealed numerically greater tracer-positive area and fluorescence intensity in MV mice, although these differences did not reach statistical significance (**Fig. 5g and h**).

Together, these awake clearance measurements provide an anesthesia-independent complement to the CSF tracer influx experiments and demonstrate that mechanical ventilation impairs brain-to-blood tracer clearance after early recovery.

### Mechanical ventilation elicits a brain-predominant cytokine response

To determine whether mechanical ventilation-associated alterations in glymphatic transport co-occurred with a brain-predominant inflammatory response, we profiled cytokines in brain tissue and serum 6 h after the 2 h of MV under isoflurane plus i.p. fentanyl (Fig. 6a and Supplementary Fig. 4a). Anti-inflammatory cytokines (IL-4, IL-10, IL-13) and pro-inflammatory cytokines/chemokines (IL-17A, IL-12p70, GM-CSF, TNF-α, IL-2, IL-1β, IL-5, IFN-γ, IL-1α, IL-6, MCP-2, LIX, KC, and MCP-1) were quantified in both brain tissue and serum (Fig. 6b and Supplementary Fig. 4b). MV was associated with increased levels of several inflammatory mediators in brain tissue, with significant increases in TNF-α, IL-5, KC, and MCP-1 in brain tissue (**Fig. 6c**). By contrast, after 2 h of NC isoflurane plus i.p. fentanyl exposure followed by 6 h of recovery, NC mice did not show significant alterations in the measured cytokine/chemokine levels in brain tissue relative to Control mice (Fig. 6d–f), indicating that anesthetic and opioid exposure without mechanical ventilation was not sufficient to reproduce the brain inflammatory response. In serum, cytokine changes after MV were limited, with IL-1α being the only significantly increased pro-inflammatory cytokine (Supplementary Fig. 4c).

**Figure 6.**
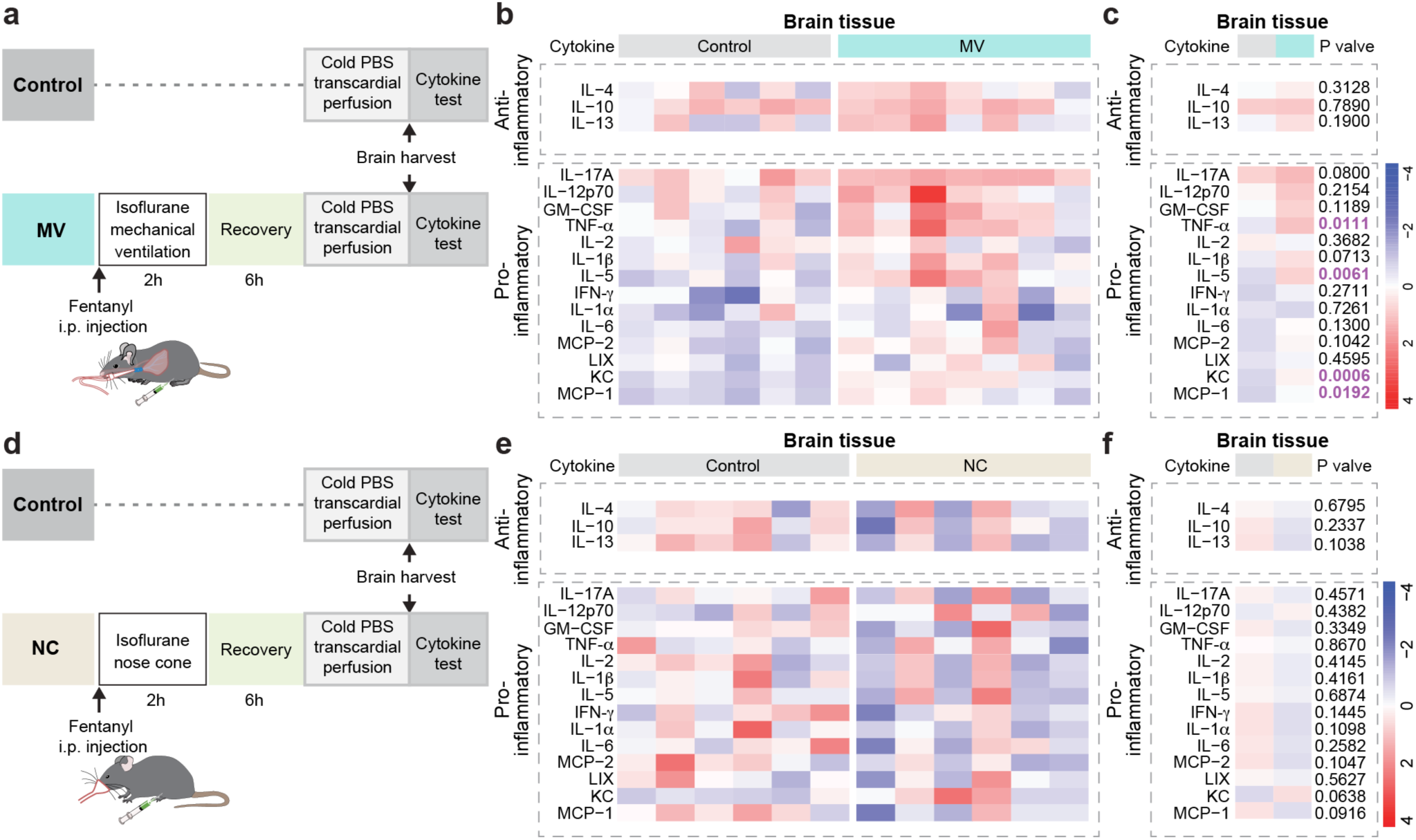
Mechanical ventilation elevates cytokine levels in brain tissue. **(a)** Assay schematic. Mice underwent 2 h of mechanical ventilation (MV) under 2.5% isoflurane anesthesia with i.p. fentanyl, followed by 6 h of recovery. Brains were then harvested for cytokine analysis. **(b)** Heatmap of cytokine levels in brain tissue from Control and MV mice. Columns denote individual mice; rows denote measured cytokines, including anti-inflammatory cytokines (IL-4, IL-10, IL-13) and pro-inflammatory cytokine /chemokines (IL-17A, IL-12p70, GM-CSF, TNF-α, IL-2, IL-1β, IL-5, IFN-γ, IL-1α, IL-6, MCP-2, LIX, KC, MCP-1). Values are scaled per cytokine to visualize relative changes (blue = low, red = high). **(c)** Group summaries of cytokine levels in Control and MV mice. Data are presented as mean ± SEM. Statistical significance was determined by two-tailed unpaired t-test. n = 6 Control mice, n = 7 MV mice. **(d)** Assay schematic. Mice received 2 h of 2.5% isoflurane anesthesia via nose cone (NC) with i.p. fentanyl, followed by 6 h of recovery. Brains were then harvested for cytokine analysis. **(e)** Heatmap of cytokine levels in brain tissue from Control and NC mice. Columns denote individual mice; rows denote measured cytokines, including anti-inflammatory cytokines (IL-4, IL-10, IL-13) and pro-inflammatory cytokines/chemokines (IL-17A, IL-12p70, GM-CSF, TNF-α, IL-2, IL-1β, IL-5, IFN-γ, IL-1α, IL-6, MCP-2, LIX, KC, MCP-1). Values are scaled per cytokine to visualize relative changes (blue = low, red = high). **(f)** Group summaries of cytokine levels in Control and NC mice. Data are presented as mean ± SEM. Statistical significance was determined by two-tailed unpaired t-test. n = 6 mice per group.

Finally, MV, but not matched NC exposure, elicited a brain-predominant inflammatory response, whereas serum cytokine changes remained limited. Together, these findings show that MV is associated with increased accumulation of inflammatory mediators in brain with only limited changes detected in serum under the conditions tested.

## DISCUSSION

Mechanical ventilation is required for ∼20–40% of ICU admissions, and in some cohorts more than half of ventilated patients receive respiratory support within the first 24 h of admission^33^. Delirium in this setting is likely to arise from the combined effects of critical illness, sedative exposure, analgesia, and ventilation-associated physiological stress. Indeed, greater early sedation intensity, MV, and opioid use have each been associated with increased delirium risk and poorer long-term outcomes^7,34–36^. Yet, how MV-associated physiological perturbations contribute to acute brain dysfunction beyond concomitant sedative and opioid exposure remains poorly defined. In our experimental model, we found that a brief period of MV administered with inhaled anesthesia and opioid analgesia was sufficient to induce a behavioral phenotype consistent with delirium-like dysfunction 24 h later in healthy mice. Across complementary behavioral assays, MV mice showed reduced exploratory engagement, increased avoidance-biased behavior, and altered spatial search behavior, whereas rotarod performance was preserved. This pattern makes gross motor impairment an unlikely primary explanation for the behavioral changes and is consistent with altered exploration, avoidance, and behavioral organization. Notably, matched nose-cone isoflurane exposure with opioid administration did not reproduce these abnormalities. Thus, these findings suggest that anesthetic and opioid exposure alone is insufficient to produce persistent cognitive disturbances. Rather, they indicate that physiological alterations associated with mechanical ventilation may disrupt brain homeostasis and contribute to delirium pathogenesis.

A growing body of clinical and translational evidence suggests that many conditions that increase delirium risk, including aging, neurodegenerative disease, sleep disruption, systemic inflammation, sedative and opioid exposure, and critical illness, also converge on impaired glymphatic transport^1,17,21,35,37–39^. This convergence raises the possibility that impaired glymphatic transport represents a shared physiological vulnerability through which diverse ICU-associated exposures may disturb brain homeostasis, potentially altering the handling of metabolic by-products and inflammatory mediators relevant to cognitive function. Consistent with this broader view, traditional antipsychotic strategies, including haloperidol, have shown limited efficacy in delirium management^40^. Although this therapeutic limitation does not define the underlying mechanism, it reinforces the concept that delirium is unlikely to reflect a primary psychotic process alone^40^ and may instead arise from a more fundamental disturbance of brain homeostasis. Our model - mechanical ventilation with inhaled anesthesia and intraperitoneal opioid administration, designed to replicate a common ICU scenario - acutely reduced glymphatic influx, altered downstream CSF efflux, was associated with reduced brain-to-blood tracer clearance, and a brain-predominant cytokine response. These abnormalities co-occurred with delirium-like behavioral changes and together define a mechanistically tractable physiological phenotype following MV (**Fig. 7**). Conversely, matched nose-cone isoflurane with fentanyl largely preserved glymphatic tracer transport and did not reproduce the behavioral phenotype, indicating that anesthetic and opioid exposure alone was insufficient to account for the MV-associated changes. Compared with matched NC mice, MV was associated with a greater increase in ICP, reduced glymphatic influx and tracer clearance, brain-predominant inflammation, and delirium-like behavioral changes. These findings suggest that the MV procedure, including intubation and positive-pressure ventilation, may contribute to delirium-like acute brain dysfunction by disrupting brain-fluid transport and inflammatory homeostasis. Future studies should determine whether early glymphatic impairment predicts later behavioral severity and whether restoring glymphatic transport improves outcomes. Thus, MV-associated disruption of brain-fluid transport, coupled with brain-predominant inflammation, provides a plausible mechanistic framework for testing mechanisms underlying MV-associated delirium-like acute brain dysfunction ^27^.

**Figure 7.**
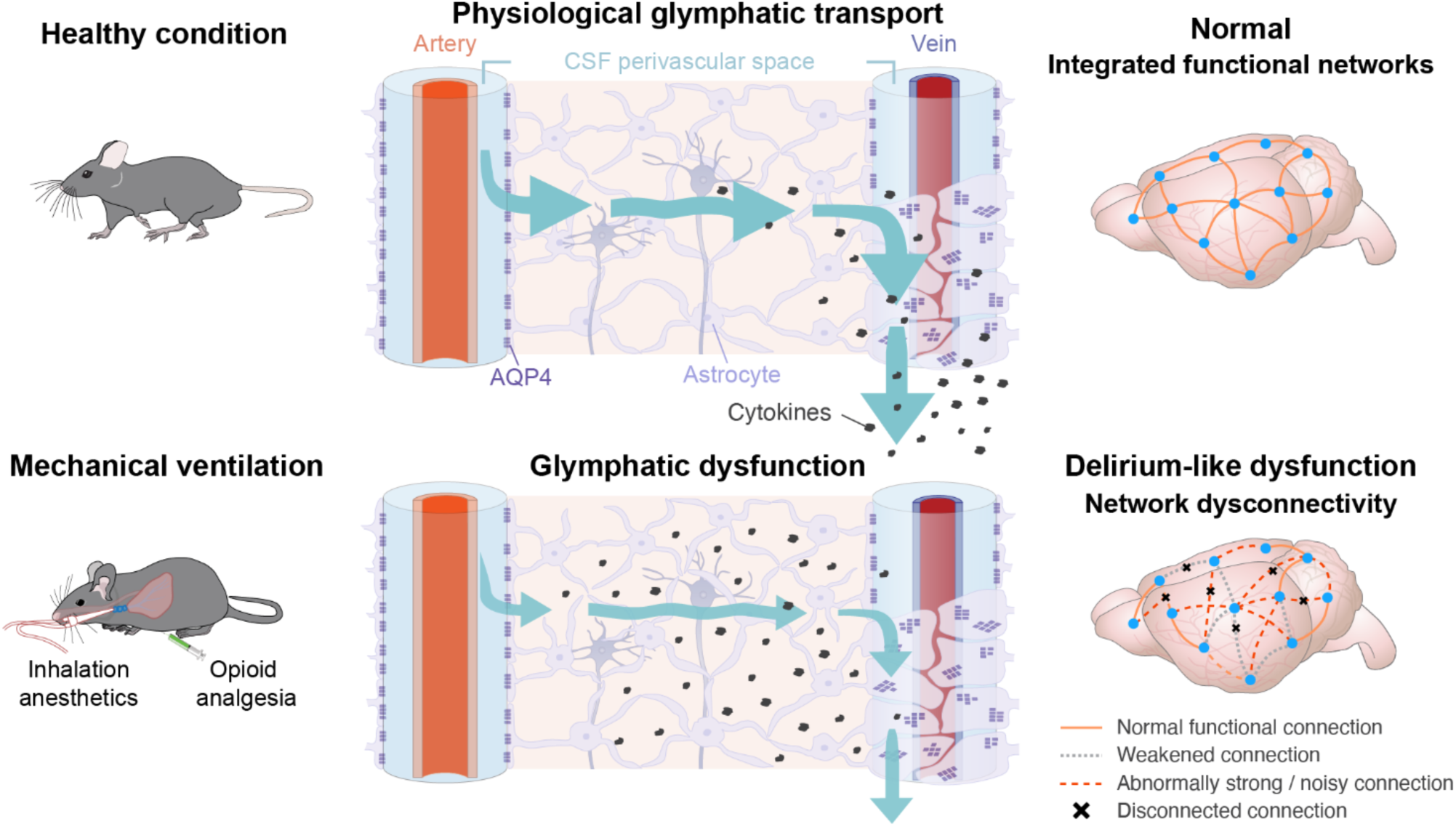
Proposed model linking mechanical ventilation-induced suppression of glymphatic transport to delirium-like behavior. Under physiological conditions, cerebrospinal fluid (CSF) enters along peri-arterial spaces, exchanges with interstitial fluid (ISF) in the parenchyma, and exits along peri-venous routes toward downstream lymphatic drainage, thereby supporting the clearance of inflammatory mediators and metabolic waste from the brain. During mechanical ventilation under inhaled anesthesia and opioid analgesia, glymphatic transport is impaired, leading to reduced CSF–ISF exchange and brain-to-peripheral clearance. This impaired clearance may promote the accumulation of cytokines and other potentially neuroactive solutes, which could disrupt neural homeostasis and functional network integration, contributing to delirium-like behavioral abnormalities.

Beyond the reduction in glymphatic tracer influx, our data suggest that MV is associated with time-dependent changes in CSF tracer distribution across downstream efflux compartments. During ventilation, tracer distribution was broadly reduced across intracranial and spinal compartments, with a concurrent increase in tracer accumulation in the scLNs. In early recovery, scLN tracer accumulation returned to levels comparable to those in Control and NC mice, whereas persistent reduction in cranial-base tracer signal was accompanied by increased tracer accumulation in the spinal cord and dcLNs. This evolving pattern suggests that MV may alter the distribution of tracer across CSF outflow-associated compartments, including cranial-base, spinal, and cervical lymphatic compartments. The persistent reduction in tracer signal within ON- and TGN-associated cranial-base regions is notable because these sites represent anatomically distinct CSF-facing perineural interfaces. The olfactory/cribriform pathway is an established route for CSF egress to nasal lymphatics^41–43^, whereas recent work has shown that CSF solutes can also access the trigeminal ganglion through the trigeminal nerve root^44^, enabling direct CSF-mediated communication between the central and peripheral nervous systems. Accordingly, the sustained reduction in tracer signal at both ON- and TGN-associated cranial-base regions after MV suggests perturbation of a broader cranial-base CSF interface, with potential implications for both olfactory-associated lymphatic egress and trigeminal access to CSF-associated solutes.

Acute elevation in ICP during MV provides a plausible physiological context for the evolving pattern of CSF efflux observed in our study. Importantly, end-tidal CO₂ was maintained within the physiological range throughout MV, making overt CO₂ retention an unlikely primary explanation for the increased ICP. Recent work showed that increased mean ICP enhances nasal-to-scLN tracer drainage without a corresponding increase in dcLN drainage ^31^. Consistent with this observation, elevated ICP during MV coincided with increased tracer accumulation in the scLNs, despite reduced tracer signal at ON- and TGN-associated cranial-base interfaces. Thus, the reduced cranial-base signal should not be interpreted simply as reduced cervical lymphatic drainage; instead, it may reflect altered tracer transit across these cranial-base interfaces together with greater tracer accumulation in the scLNs. This pattern evolved during early recovery: although scLN tracer accumulation returned to control-like levels, persistent reductions in cranial-base tracer signal were accompanied by increased tracer accumulation in the spinal cord and dcLNs. Experimental elevation of ICP has been shown to impair glymphatic influx ^30^. Similarly, in experimental cortical ischemia, ICP elevation was associated with faster CSF transit through the spinal subarachnoid space when cranial clearance was impaired ^45^. Together, these observations are consistent with the possibility that transient ICP elevation during MV is associated with, and may contribute to, a time-dependent rerouting of CSF transport across cranial base, cervical lymphatic, and spinal pathways. The recent identification of leptomeningeal arterial–venous overlap zones that permit CSF-associated fluid and macromolecule shunting provides a potential anatomical substrate for such redistribution ^46^. Future studies should determine whether preventing or attenuating MV-associated ICP elevations is associated with preservation of glymphatic influx and normalization of CSF tracer distribution across cranial base, cervical lymphatic, and spinal pathways.

Pro-inflammatory cytokines are strongly implicated in delirium pathogenesis because inflammatory signaling can disrupt neural processes underlying attention and cognition ^47–49^, and clinical inflammatory stressors such as sepsis and surgery further increase delirium risk ^50^. We found that mechanical ventilation in combination with inhalation anesthesia and opioid analgesia was sufficient to elicit a brain-predominant cytokine response within 6 h in otherwise healthy mice, without imposed systemic disease or surgical injury. TNF-α, which has well-established neuroinflammatory and cognitive effects ^51^, and MCP-1, a chemokine involved in monocyte recruitment and microglial activation ^52^, were both elevated after MV. IL-5 and KC were also increased, indicating a broader early cytokine and chemokine response following MV. The absence of a comparable cytokine response in matched mice that were not mechanically ventilated further implicates mechanical ventilation–associated physiology, rather than anesthetic and opioid exposure alone, as a driver of this brain-predominant inflammatory phenotype. The co-occurrence of reduced glymphatic tracer transport and increased levels of selected brain cytokines is consistent with the possibility that glymphatic transport normally contributes to the clearance or washout of cytokines and other inflammatory mediators from the CNS. Reduced glymphatic transport after MV could therefore limit the removal of such mediators and contribute to their local persistence. Although the temporal pattern and established biological role of glymphatic transport support the possibility that reduced glymphatic clearance contributes to the persistence of inflammatory mediators in the brain after MV, targeted manipulation of glymphatic transport will be needed to establish the magnitude of this contribution. Notably, our data showed that cytokine changes were brain-predominant, with only limited changes detected in serum. This brain-predominant pattern is consistent with CNS immune compartmentalization, in which central and systemic cytokine milieus can diverge ^53^; indeed, hippocampal cytokine changes have been reported in the absence of corresponding serum changes ^54^. Taken together, our findings show that MV-associated alterations in glymphatic tracer transport co-occurred with a CNS-predominant cytokine response, providing a framework for future studies to test the direction and mechanisms of this relationship.

Animal models of delirium are essential for advancing the neurobiology of the syndrome and for testing treatments. Many current models combine a predisposing vulnerability (e.g., advanced age or neurodegeneration) with a clinically relevant precipitant (e.g., infection, anesthesia, surgery) ^28,55,56^. Although these models in part replicate that delirium is multifactorial, combining multiple vulnerabilities and precipitating factors can complicate causal attribution and make it difficult to distinguish delirium-like behaviors from the effects of systemic illness, surgery, or underlying disease ^28^. In contrast to inflammation-based paradigms that rely on lipopolysaccharide (LPS) or surgery (such as laparotomy or tibial fracture), both of which introduce systemic confounds, our model uses otherwise healthy mice and a defined, clinically relevant exposure paradigm consisting of mechanical ventilation with inhalational anesthesia and adjunct opioid analgesia. This approach models key features of a routine peri-operative/ICU exposure while avoiding the deliberate induction of severe systemic inflammation, hypoglycemia, or global ischemia, thereby facilitating interpretation of the observed brain-predominant cytokine response. The model is feasible in adult mice and provides clear physiological and behavioral readouts, enabling rigorous evaluation of preventive and therapeutic strategies and supporting predictive validity. More broadly, this model provides a tractable platform for testing alternative hypotheses about delirium pathogenesis.

In conclusion, mechanical ventilation administered under inhaled anesthesia and opioid analgesia was associated with reduced glymphatic tracer influx, altered tracer distribution across downstream CSF outflow-associated compartments, reduced brain-to-blood tracer clearance, elevated ICP, and a brain-predominant cytokine response. These abnormalities co-occurred with delirium-like behavioral dysfunction in otherwise healthy mice. Together, these findings define a mechanistically tractable model in which relationships among MV-associated physiological stress, brain-fluid transport, inflammation, and delirium-like behavioral dysfunction can be directly tested. Therapeutic strategies aimed at modulating brain-fluid transport, CSF outflow-associated pathways, or adverse ICP elevations warrant direct evaluation as approaches to mitigate delirium-related brain dysfunction in critical care.

## MATERIALS AND METHODS

### Animals

All animal procedures were approved by the University Committee on Animal Resources at the University of Rochester and conformed to the National Institutes of Health (NIH) Guide for the Care and Use of Laboratory Animals and the ARRIVE guidelines. Male and female C57BL/6 mice (Charles River Laboratories, Wilmington, MA) were housed under controlled conditions (temperature 70–74 °F, humidity 30–70%) on a 12:12 h light–dark cycle until they reached the required ages for experimentation. Four-month-old mice were used for intracisternal infusion studies; ten-month-old mice were used for clearance assays, behavioral assessments, intracranial pressure measurements, and cytokine analyses. Group sizes were determined based on prior experience, technical feasibility, and published reports. Animals were randomly assigned to experimental groups.

### Mechanical Ventilation

Animals were induced with 4% isoflurane in oxygen and orotracheally intubated using a 20G catheter. After intubation, mice were ventilated on a small-animal ventilator (RoVent Jr., Kent Scientific) with the following settings: tidal volume 6.65 mL·kg⁻¹, respiratory rate 116–139 breaths·min⁻¹ initially adjusted for body weight and subsequently fine-tuned based on end-tidal CO₂ values, and target peak inspiratory pressure (PIP) 7 cm H₂O. Anesthesia was maintained at 2.5% isoflurane via the ventilator. Animals were placed on an electronic heating pad (37°C; Stoelting Rodent Warmer). End-tidal CO₂ was monitored with CapnoScan (Kent Scientific), and respiratory rate was adjusted as needed to maintain end-tidal CO₂ within the normal physiological range of 30–45 mmHg. Electrocardiogram and hind-limb pulse oximetry using a sensor affixed to the shaved hindlimb were monitored with a small-animal physiological monitoring system (Harvard Apparatus).

### Drugs

Immediately following start of mechanical ventilation, animals received fentanyl (2.5% w/v in Saline, 100 μg/kg) intraperitoneally. Ketamine/xylazine was administered 100/10 mg·kg⁻¹ i.p. The isoflurane concentration delivered by nose cone was 2.5%.

### Behavioral Tests

All behavioral experiments were performed 24h after completion of the mechanical ventilation procedure.

### Open Field Test

Each animal was placed in the center of a rectangular arena (45 × 60 cm) and allowed to explore freely for 10 min.

### Barnes maze

Animals received 3 training sessions per day for 2 days prior to mechanical ventilation. In each training session, the animal was placed in a dark box on Barnes maze platform for 15s and allowed to explore in Barnes maze with spatial cues for 2 min. After successfully entering the escape box within 2 min or being guided into the escape box after 2 min, the animal was allowed to stay in the escape box for 1 min. The interval between training sessions for each animal was 20-25 min. During the test, the animal was placed in a dark box on Barnes maze platform for 15s and allowed to explore in Barnes maze for 2 min.

### Light-dark box test

The light dark box apparatus (l × w × h = 45 × 27 × 30 cm) consists of one third of dark compartment and two thirds of light compartment^57^. Two compartments are separated by a gate (7.5 × 7.5 cm). At the beginning of the test, the animal is placed at the corner of the dark compartment and is allowed to freely explore in the light dark box for 5 minutes. The light compartment exploration time and numbers of visit are recorded.

### Rotarod test

Mice were placed on a continuously rotating accelerating rota-rod (7650, Ugo-Basile, Gemonio, Italy). Once mice were secure on the rotation beam at the lowest spin speed, the rotarod was switched to acceleration mode. The time from the beginning of the acceleration until the mouse fell off of the rotating beam was recorded. Mice were allowed to rotate on the beam for at most 5 minutes.

Three trails were obtained for each mouse. Mice were given at least 5 minute of rest time in home cage in between each trial. The average latency for the mice to fall off the beam and the latency for the best performance among three trials were calculated.

All behavioral experiment videos were recorded with a surveillance system (Reolink). All behavioral recordings were analyzed using AnyMaze (Stoelting) by an experimenter blinded to group allocation.

### Intracisternal Cannulation

The mice were anesthetized with ketamine/xylazine intraperitoneally and positioned in a stereotaxic frame. A midline incision was made to expose the skin above skull and back neck. Underlying connective tissue and cervical muscles were carefully separated along the rostro caudal midline to expose the cisterna magna (CM). A PE-10 tube cannulated with 30-gauge needle was preloaded with a fluorescence tracer (0.5% w/v in aCSF, Albumin from bovine serum, Alexa Fluor® 647 conjugate, 66kDa, Invitrogen) and connected to a syringe pump (Harvard Apparatus). The needle was inserted into CM and secured with cyanoacrylate adhesive (Krazy Glue). A metal head plate implanted over skull was secured by a head bar before transcranial imaging.

### Transcranial Imaging

Following intracisternal cannulation, animals were positioned under an epifluorescence microscope (MVX10, Olympus) with a pE-400 LED light source (CoolLED) and an ORCA Flash 4.0 digital camera (Hamamatsu) for imaging. Fluorescence tracer was infused into the CM with a rate of 2μL/min for 5 min with a syringe pump (Harvard Apparatus). Transcranial imaging started simultaneously with tracer infusion, and images were acquired with a sample rate of 1 image per min for 30 min from both far-red (647nm) and green (450nm) emission channels.

### Tissue Processing

Following transcranial imaging, mice received 100 μL of WGA-488 (1 mg/mL) via the femoral vein immediately before decapitation. The superficial and deep cervical lymph nodes, whole brain, skull, and spinal cord were collected and fixed in 4% paraformaldehyde (PFA) at 4 °C for 24 h. Fixed brains were sectioned coronally at 100 μm thickness using a vibratome (VT1200 S, Leica Biosystems). The tissues were imaged using either an epifluorescence microscope (MVX10, Olympus) or a confocal microscope (FV3000, Olympus; FV31S software) equipped with a ×10 objective lens (NA 0.40) and EGFP and Cy5 filter sets.

For the brain-to-blood clearance assay, brains were collected after fiber-photometry recording, fixed in 4% PFA at 4 °C for 24 h, and sectioned coronally at 100 μm using a vibratome (VT1200 S, Leica Biosystems). Residual Cy5-biotin in ex vivo brain sections was then evaluated using an epifluorescence microscope (MVX10, Olympus).

### Glymphatic Clearance Assay

Twenty-four hours before fiber-photometry recording, mice were induced with 4% isoflurane and maintained at 2.5% isoflurane after placement in a stereotaxic frame. Pre-emptive analgesia was provided with carprofen (5 mg·kg⁻¹, s.c.) and bupivacaine (2.5 mg·mL⁻¹, local infiltration) at the surgical site prior to implantation. After skull exposure, a burr hole was drilled at AP +0.3 mm, ML −2.5 mm (relative to bregma) using a microdrill (Harvard Apparatus). A guide infusion cannula (4.5-mm projection, dust cap; RWD Life Science) was implanted into the striatum at DV −3.8 mm. In addition, a 200-μm core, 1.5-mm projection fiber-optic cannula (RWD Life Science) was positioned over the confluence of the superior sagittal sinus (SSS) and inferior cerebral vein (ICV). Both the infusion and fiber-optic cannulae were secured with adhesive cement (C&B Metabond, Parkell). Animals were then habituated in the recording chamber prior to fiber-photometry acquisition.

Following 2 h of mechanical ventilation, animals recovered awake for 1 h in the recording chamber. A PE-10 line coupled to a 4.5-mm projection injector (RWD Life Science) was preloaded with mineral oil (Sigma-Aldrich) and Cy5-conjugated biotin (1% w/v in aCSF; Vector Labs), mounted on a micro-syringe pump (Pump 11 Elite, Harvard Apparatus), and inserted into the guide infusion cannula after removal of the dust cap. A 2-m optic fiber (200-μm core, NA 0.22, FC/PC with Ø1.25-mm ceramic ferrule; RWD Life Science) was connected to a fluorescence mini-cube (Doric) and a fiber-photometry system (RZ10-X Lux-I/O with 405/465/635-nm excitation LEDs and LxPS1 photodetectors; TDT) and secured to the implanted fiber-optic cannula. Fifteen minutes before recording, topical bupivacaine (2.5 mg·mL⁻¹) was applied to the right eye, followed by retro-orbital streptavidin (5% w/v in saline; Neuromics) into the venous sinus. Fiber-photometry acquisition was initiated concurrently with the infusion (0.1 μL·min⁻¹ for 10 min). Recordings continued for 2 h with animals undisturbed. 635-nm excitation was delivered through the optic fiber to excite the Cy5–biotin–streptavidin complex. At the end of recording, animals were decapitated for tissue collection.

For data analysis for glymphatic clearance, CY5 signal over the confluence of SSS and ICV was calculated using 635 nm signal in recording and normalized with a moving baseline in MATLAB.

### ICP Measurement

Animals were induced with 4% ISO and anesthetization was maintained with 2.5% ISO after the animals were placed onto a stereotaxic frame. After skull exposure, a burr hole was drilled (AP: −0.3, ML: −1.0 mm) using Micro Drill (Harvard Apparatus). A PE10 tube cannulated to a 30-gauge dental needle and preloaded with aCSF was lowered into the lateral ventricle (DV: −2.0 mm). The dental needle was secured with a mixture of cyanoacrylate adhesive and dental cement powder (Stoelting).

The implanted PE-10 line was connected to a pressure transducer (WPI) prior to recording. The transducer output was routed to a transducer amplifier (TMB4M, WPI) and a digitizer (Axon Digidata 1550B, Molecular Devices) to convert analog signals to digital data. AxoScope 10 (Molecular Devices) was used for acquisition of ICP. At recording onset, animals underwent either 2 h of mechanical ventilation or 2 h of isoflurane anesthesia delivered via nose cone, according to group assignment, and were maintained on a 37 °C heating pad throughout. After the 2-h procedure, animals were removed from the ventilator or nose cone and allowed to recover for 1 h, after which ICP recording was terminated.

### Cytokine Test

Six hours after the 2-h mechanical ventilation procedure, animals were re-anesthetized with ketamine/xylazine (i.p.). Blood was collected from the abdominal aorta using a 27-gauge needle and allowed to clot at room temperature for 30 min; samples were then centrifuged at 3,000 rpm for 10 min at 4 °C to isolate serum. Animals were subsequently transcardially perfused with ice-cold PBS (4 °C) to remove residual blood. Whole brains were harvested and homogenized on ice in RIPA lysis buffer (25 mM Tris·HCl pH 7.6, 150 mM NaCl, 1% NP-40, 1% sodium deoxycholate, 0.1% sodium dodecyl sulphate (SDS) (Thermo Fisher) containing a protease inhibitor cocktail (1:100 ratio, Sigma-aldrich, Burlington, MA). Homogenates were sonicated (SFX150, Branson; 40 kHz, 10 s) and centrifuged at 13,000 rpm for 20 min at 4 °C to collect supernatants. Protein concentrations were determined by BCA assay (Thermofisher, Waltham, MA). Serum and brain homogenates that met the platform’s concentration requirements were shipped to Eve Technologies Company for cytokine profiling using the Mouse High Sensitivity T-Cell 18-Plex Discovery Assay® (MDHSTC18).

### Statistical Analysis

All quantification was performed in Fiji or MATLAB, and statistical analyses were conducted in GraphPad Prism 10. Representative images were chosen from the full dataset and are identified in the figure legends. Statistical tests included two-tailed unpaired t-test or one-way analysis of variance (ANOVA) post hoc Tukey’s test, and two-way ANOVA with Sidak’s multiple-comparisons post hoc test. A threshold of P < 0.05 was considered statistically significant. Data are reported as mean ± SEM. Sample sizes were not predetermined by power analysis but are comparable to those in prior publications ^58,59^. No animals were excluded except in cases of anesthesia failure or technical issues during recording.

## Acknowledgments

We would like to thank Dan Xue for assistance with the illustrations.

## Author contributions

Conceptualization: T.D., G.L., M.N.; Investigation: T.D., G.L., J.W., T.T.; Methodology: T.D., G.L., J.W., T.T., S.H., E.N., E.K.; Formal analysis: T.D., J.W., T.T., G.L., A.L., S.L.; Writing-original draft: T.D., M.N., Writing-review and editing: T.D., M.N., G.L., J.W., T.T., S.H., A.L., E.N., E.K., H.C.B.

## Supplementary Figures and Legends

**Supplementary Figure 1.**
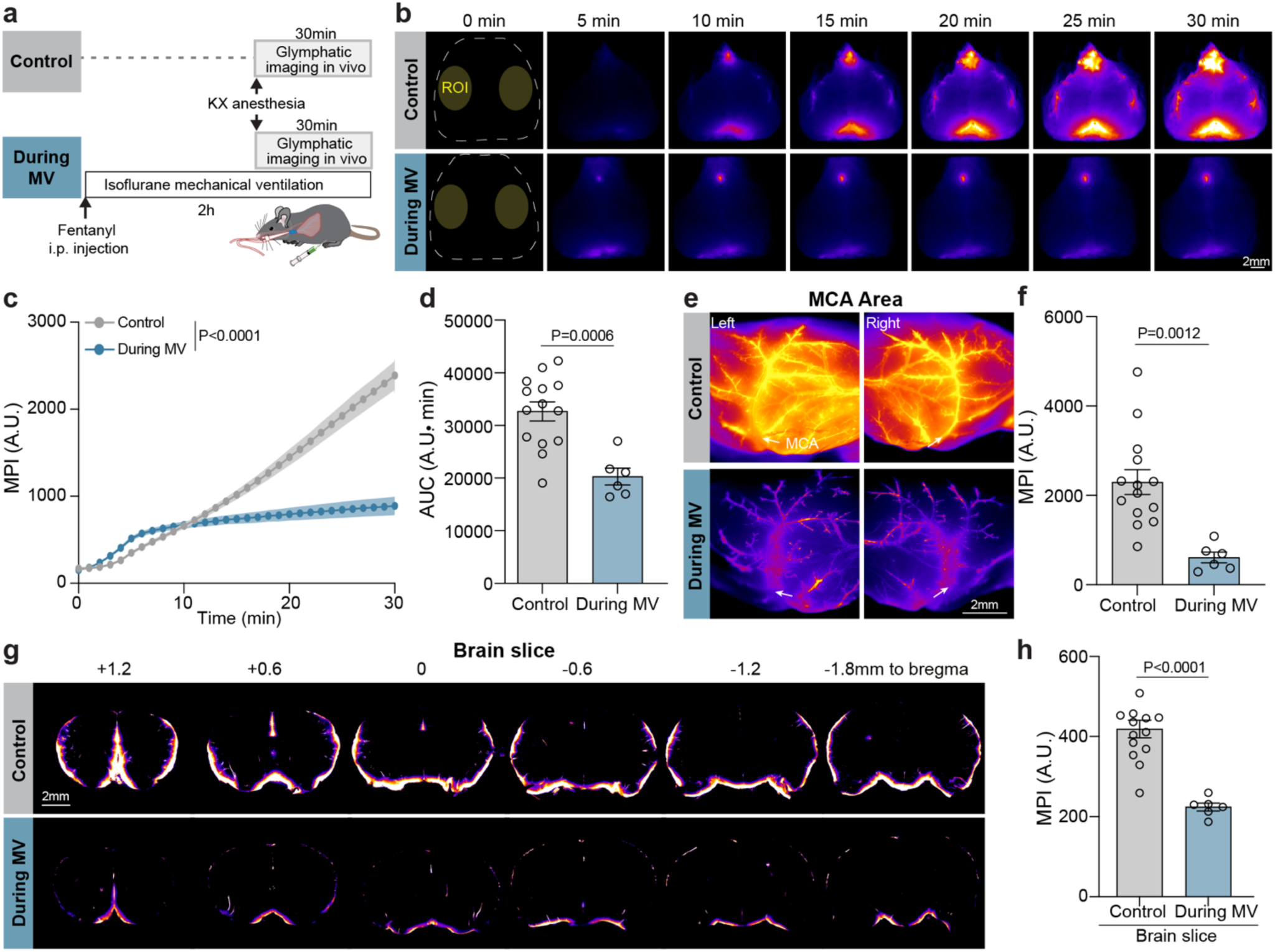
Glymphatic transport impairment is already evident during mechanical ventilation. **(a)** Experimental timeline for assessing glymphatic tracer influx during ongoing mechanical ventilation. Control mice under ketamine/xylazine (KX) anesthesia received a cisterna magna (CM) injection of BSA–Alexa Fluor 647 (BSA-647), followed by 30 min of transcranial imaging. Mice in the During MV group underwent 2 h of mechanical ventilation under 2.5% isoflurane anesthesia after intraperitoneal (i.p.) fentanyl injection. At 1.5 h after the start of MV, mice received KX anesthesia for CM injection of BSA-647, followed by 30 min of transcranial imaging during ongoing MV. **(b)** Representative transcranial fluorescence images showing time-lapse intracisternal tracer influx in Control and During MV mice. Scale bar, 2 mm. **(c)** Quantification of mean pixel intensity (MPI) of tracer influx over 0–30 min within the middle cerebral artery (MCA) territory. **(d)** Area under the curve (AUC) of 30-min BSA-647 influx shown in panel c. **(e)** Representative high-resolution ex vivo fluorescence images of intact whole brains showing tracer distribution in the MCA territory. **(f)** Quantification of ex vivo MPI in the MCA territory. **(g)** Representative ex vivo fluorescence images of coronal sections from anterior to posterior. Scale bar: 2 mm. **(h)** Quantification of MPI in ex vivo coronal brain sections. The Control data are the same dataset shown in Fig. 2 and were included here as a shared reference group for comparison with the During MV group. Data are presented as mean ± SEM. Statistical significance was determined by two-way repeated-measures ANOVA followed by Sidak’s multiple-comparisons test for panel c and by two-tailed unpaired t-test for panels d, f, and h. n = 14 Control mice and n = 6 During MV mice. ns, not significant.

**Supplementary Figure 2.**
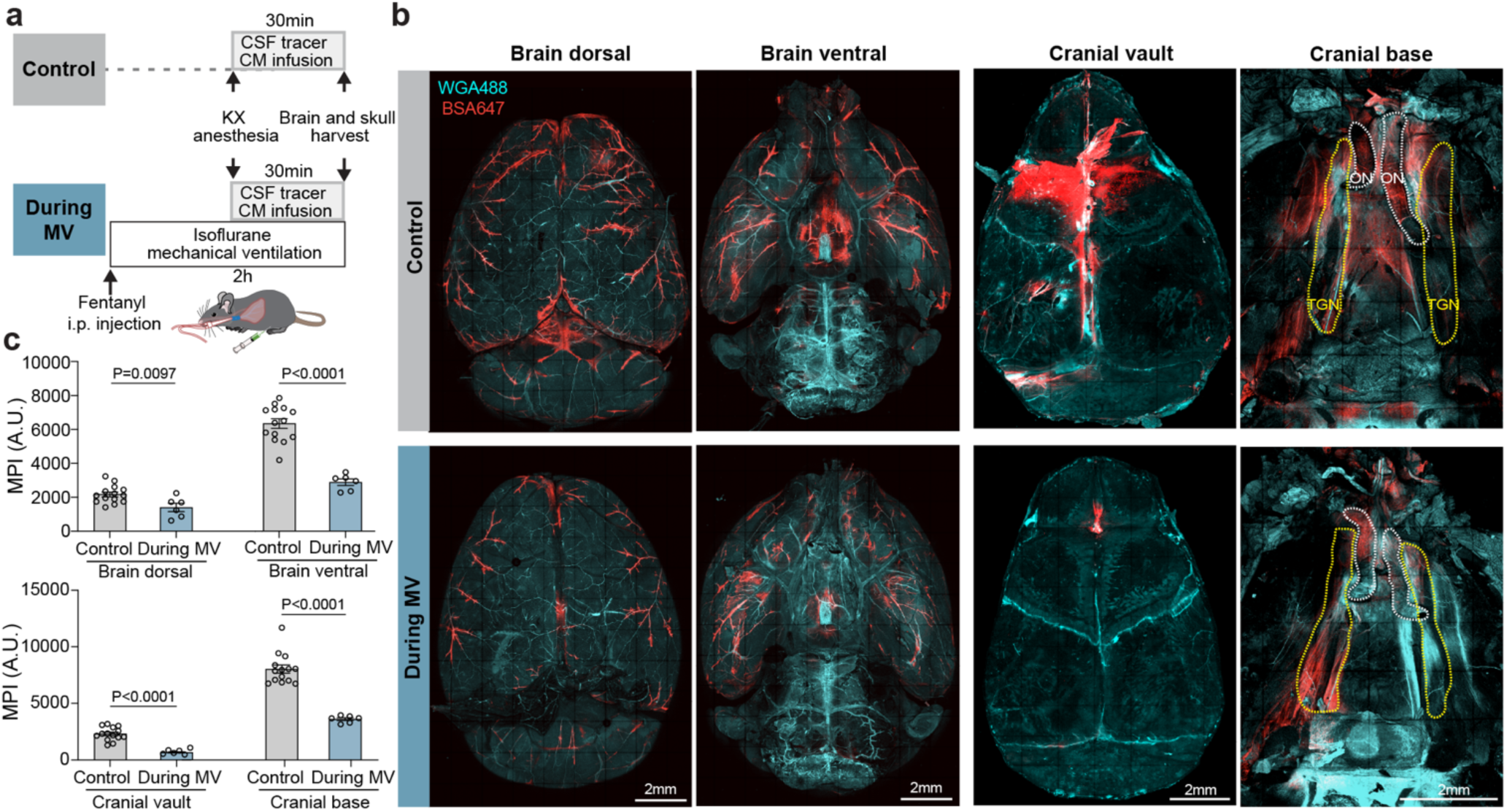
Mechanical ventilation acutely alters intracranial CSF tracer distribution during ventilation. **(a)** Experimental timeline for ex vivo assessment of intracisternally delivered tracer distribution during ongoing mechanical ventilation. Control mice under ketamine/xylazine (KX) anesthesia received a cisterna magna (CM) injection of BSA–Alexa Fluor 647 (BSA-647), followed by a 30-min influx period. Mice in the During MV group underwent mechanical ventilation under 2.5% isoflurane anesthesia after intraperitoneal (i.p.) fentanyl injection. At 1.5 h after the start of a 2 h MV bout, mice received KX anesthesia for CM injection of BSA-647, followed by a 30-min influx period during ongoing MV. Whole brains and skulls were harvested from Control and During MV mice and imaged ex vivo by fluorescence microscopy. **(b)** Representative dorsal and ventral whole-brain fluorescence images, together with cranial vault and cranial base fluorescence images, from Control and During MV mice. Scale bar, 2 mm. **(c)** Quantification of mean pixel intensity (MPI) in the dorsal brain, ventral brain, cranial vault, and cranial-base regions associated with the olfactory nerve (ON) and trigeminal nerve (TGN). The Control data are the same dataset shown in Fig. 3 and were included here as a shared reference group for comparison with the During MV group. Data are presented as mean ± SEM. Statistical significance was determined by two-tailed unpaired t-test. n = 14 Control mice and n = 6 During MV mice.

**Supplementary Figure 3.**
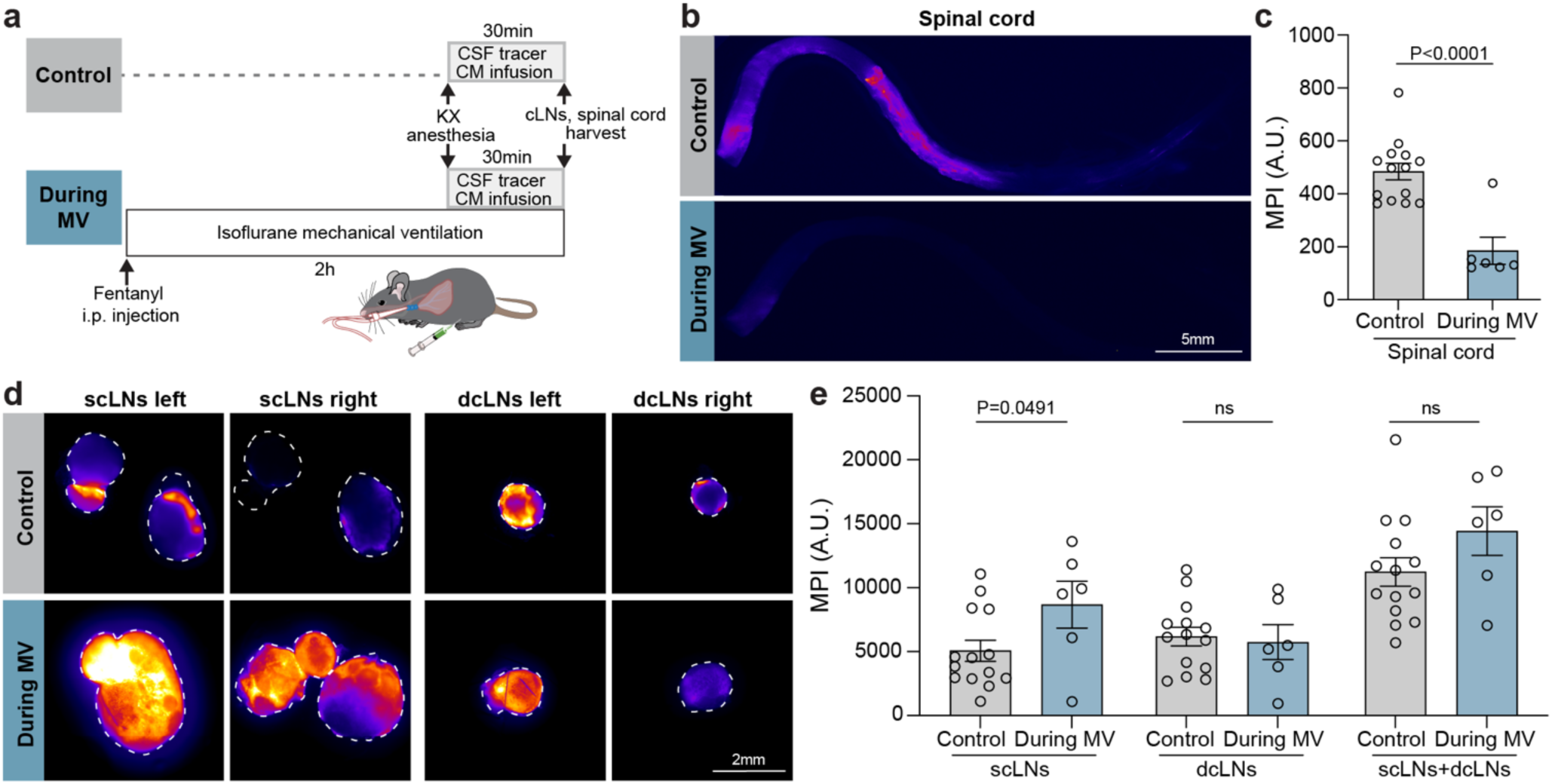
CSF efflux paths are altered during mechanical ventilation. **(a)** Experimental timeline for ex vivo assessment of intracisternally delivered tracer distribution. Following cisterna magna (CM) injection of BSA–Alexa Fluor 647 (BSA-647) and a 30-min influx period, as described in Supplementary Fig. 2a, tissues including the spinal cord, superficial cervical lymph nodes (scLNs), and deep cervical lymph nodes (dcLNs) were harvested from Control, During mechanical ventilation (MV) mice and imaged ex vivo by fluorescence microscopy. **(b)** Representative fluorescence images of the spinal cord. Scale bar, 5 mm. **(c)** Bar graph shows quantification of spinal cord MPI. **(d)** Representative fluorescence images of scLNs and dcLNs from Controland During MV mice. Scale bar, 2 mm. **(e)** Bar graphs show quantification of MPI in scLNs, dcLNs, and combined scLNs + dcLNs. The Control data are the same dataset shown in Fig. 4 and are included here as a shared reference group for comparison with the During-MV group. Data are presented as mean ± SEM. Statistical significance was determined by two-tailed unpaired t-test were performed. n = 14 Control mice and n = 6 During MV mice. ns, not significant.

**Supplementary Figure 4.**
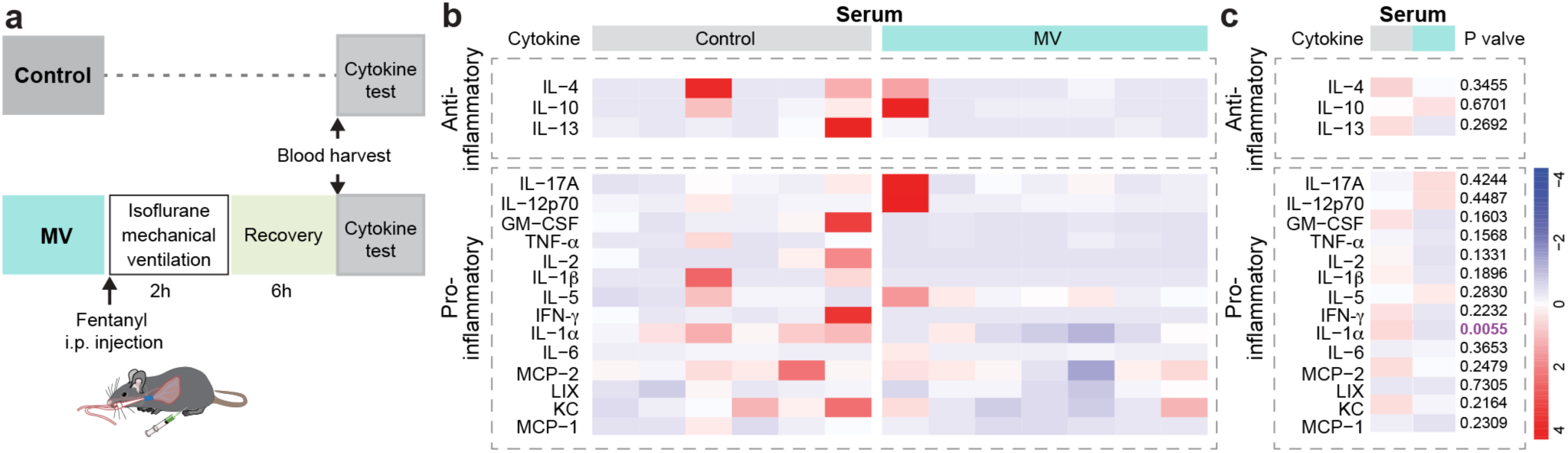
Mechanical ventilation does not broadly elevate serum cytokine levels. **(a)** Assay schematic. Mice underwent 2 h of mechanical ventilation (MV) with 2.5% isoflurane plus i.p. fentanyl, followed by 6 h of recovery. Blood was collected and centrifuged. The serum was then harvested for cytokine analysis. Serum samples were collected from the same mice used for brain cytokine analysis in Fig. 6a–c. **(b)** Heatmap of cytokine levels in serum from Control and MV mice. Columns denote individual mice; rows denote measured cytokines, including anti-inflammatory cytokines (IL-4, IL-10, IL-13) and pro-inflammatory cytokine /chemokines (IL-17A, IL-12p70, GM-CSF, TNF-α, IL-2, IL-1β, IL-5, IFN-γ, IL-1α, IL-6, MCP-2, LIX, KC, MCP-1). Values are scaled per cytokine to visualize relative changes (blue = low, red = high). **(c)** Group summaries of serum cytokine/chemokine levels in Control and MV mice. Data are presented as mean ± SEM. Statistical significance was determined by two-tailed unpaired t-test. n = 6 Control mice, n = 7 MV mice.

**Video 1. Glymphatic influx after 1 h recovery from mechanical ventilation.**

Control mice under ketamine/xylazine (KX) anesthesia received a cisterna magna (CM) injection of BSA–Alexa Fluor 647 (BSA-647), followed by 30 min of transcranial imaging. NC mice received 2 h of 2.5% isoflurane anesthesia delivered via a nose cone, with intraperitoneal (i.p.) fentanyl administered before isoflurane exposure, were allowed to recover awake for 1 h, and were then re-anesthetized with KX before receiving a CM injection of BSA-647, followed by 30 min of transcranial imaging. MV mice underwent 2 h of mechanical ventilation under 2.5% isoflurane anesthesia, with i.p. fentanyl administered before ventilation, recovered awake for 1 h, and were then re-anesthetized with KX before receiving a CM injection of BSA-647, followed by 30 min of transcranial imaging. Scale bar, 2 mm.

**Video 2. Glymphatic influx during mechanical ventilation.**

Control mice under ketamine/xylazine (KX) anesthesia received a cisterna magna (CM) injection of BSA–Alexa Fluor 647 (BSA-647), followed by 30 min of transcranial imaging. Mice in the During MV group underwent a 2 h MV bout under 2.5% isoflurane anesthesia, with i.p. fentanyl administered before ventilation. At 1.5 h after the start of MV, mice were additionally administered KX, followed by CM injection of BSA-647 and 30 min of transcranial imaging during ongoing MV. Scale bar, 2 mm.

